# Monocyte reconstitution and gut microbiota composition after hematopoietic stem cell transplantation

**DOI:** 10.1101/777268

**Authors:** Sejal Morjaria, Allen W. Zhang, Sohn Kim, Jonathan U. Peled, Simone Becattini, Eric R. Littmann, Eric. G. Pamer, Miguel-Angel Perales, Michael C. Abt

## Abstract

Monocytes are an essential cellular component of the innate immune system that support the host’s effectiveness to combat a range of infectious pathogens. Hemopoietic cell transplantation (HCT) results in transient monocyte depletion, but the factors that regulate recovery of monocyte populations are not fully understood. In this study, we investigated whether the composition of the gastrointestinal microbiota is associated with the recovery of monocyte homeostasis after HCT.

**Methods:** We performed a single-center, prospective, pilot study of 18 recipients of either autologous or allogeneic HCT. Serial blood and stool samples were collected from each patient during their HCT hospitalization. Analysis of the gut microbiota was done using 16S rRNA gene sequencing and flow cytometric analysis was used to characterize the phenotypic composition of monocyte populations.

**Results:** Dynamic fluctuations in monocyte reconstitution occurred after HCT and large differences were observed in monocyte frequency among patients over time. Recovery of absolute monocyte counts and monocyte subsets showed significant variability across the heterogeneous transplant types and conditioning intensities; no relationship to the microbiota composition was observed in this small cohort.

**Conclusion:** A relationship between the microbiota composition and monocyte homeostasis could not be firmly established in this pilot study.

## BACKGROUND

Hematopoietic cell transplantation (HCT) is a potentially curative procedure for patients with hematologic malignancies but its success has been limited by the morbidity and mortality of post-transplant infections and relapse. Impaired immune reconstitution post-HCT increases the risk of both of these complications [1-4]. Multiple factors influence the vulnerability of patients to these complications including: time since transplantation, graft source (i.e., autologous (auto-HCT) or allogeneic (allo-HCT)), and persisting myelosuppression (humoral and cell-mediated) post-HCT [2, 3]. Alterations in the microbiome have been associated with clinical outcomes of patients who have undergone HCT, including their survival. However, the mechanisms by which the gut microbiota exerts its effects, both beneficial and detrimental, have not been fully elucidated [5, 6].

In humans there are three main circulating monocyte subsets with diverse functions, classified based on their expression of CD14 and CD16 surface proteins and cytokine production. These monocyte subsets are named “classical”, “intermediate”, and “non-classical monocytes” [7-9] and typically comprise 85%, 10%, and 5% respectively of the circulating monocyte pool in a healthy individual under homeostatic conditions [10, 11]. Classical monocytes specialize in phagocytosis and produce the cytokine IL-10 [12], while intermediate monocytes have elevated surface expression of MHC class II, suggesting they have an important role in antigen presentation [13]. In contrast, non-classical monocytes secrete substantial levels of inflammatory cytokines (TNF and interleukin [IL]-1β) and are known to exert endothelial surveillance by endovascular slow patrolling when tissue damage is present [14]. These roles make monocytes a critical cell line for host defense against common post-HCT infections including *Aspergillus*, and for mitigating the risk of developing GVHD [15-22].

While it has been shown that monocyte reconstitution in the first 100 days post-HCT is associated with improved survival [23, 24], a better understanding of the mechanisms that regulate monocyte reconstitution in this setting is needed. Given the known immunomodulating properties of the microbiota [25-29], we investigated whether the composition of the intestinal microbiota is associated with monocyte recovery. Monocyte migration out of the bone marrow and into the blood circulation is driven by low level lipopolysacharide (LPS)-mediated signaling [30]. Further, basal circulating LPS levels derived from the intestinal microbiota have been detected in immunocompromised hosts [31, 32]. Therefore, we hypothesized that microbial communities of gram-negative bacteria through the production of LPS [33, 34] and commensal obligate anaerobes [35] influence monocyte maturation post-HCT.

## METHODS

### Study Patients, Specimen Collection, and Patient Tracking

We followed 18 adult recipients of auto- or allo-HCT at Memorial Sloan Kettering Cancer Center (MSKCC) from July 2015 to January 2016. There were 7 female and 11 male patients; their ages ranged from 40 to 75. Fecal samples were collected longitudinally from each patient during their transplant hospitalization using a prospective institutional fecal biospecimen collection protocol (described previously) [36]. For the majority of patients, daily collection began at the start of pre-transplant conditioning (7-10 days before hematopoietic cell infusion) and continued until discharge. The median average length of stay for patients in this cohort was 27 days. Clinical metadata for all patients, including medication administrations (i.e. chemotherapy agents, antibiotics, etc.), absolute white blood cell values (obtained from routine daily complete blood counts), and other patient characteristics, were retrieved from the electronic health record. The study protocol was approved by the institutional review board. Informed consent was obtained from all subjects prior to specimen collection.

### Transplantation Practices

Antimicrobial prophylaxis was given routinely to patients undergoing HCT. Subjects undergoing either auto- or allo-HCT were given ciprofloxacin two days prior to hematopoietic cell infusion as prophylaxis against gram-negative bacterial infections. Allo-HCT recipients were given intravenous (IV) vancomycin for prophylaxis against viridans-group streptococci [28]. Antibiotic prophylaxis against *Pneumocystis jiroveci* pneumonia was generally administered using either trimethoprim-sulfamethoxazole, aerosolized pentamidine, or atovaquone; the time at which prophylaxis was initiated (during conditioning or after engraftment, defined as an absolute neutrophil count ≥ 500 neutrophils/mm3 for three consecutive days) varied. In the event of a new fever during times of neutropenia, patients were usually started on empiric antibiotics, such as piperacillin-tazobactam, cefepime, or meropenem. Recipients of an autograft received pegfiltastrim on day +1 and recipients of an allograft received daily filgastrim starting on day+7 until engraftment to accelerate recovery from neutropenia [37].

### Sample Analysis and Defining Microbial Predictors

#### Sample Analysis

Stool DNA was extracted and purified, and the V4-V5 region of the 16S rRNA gene was amplified by polymerase chain reaction using modified universal bacterial primers [38]. Sequencing was performed using the Illumina Miseq platform[39] to obtain paired-end reads. These reads were assembled, processed, filtered for quality, and grouped into operational taxonomic units of 97% similarity using a previously described UPARSE pipeline [25]. Taxonomic assignment to species level was performed using nucleotide BLAST (Basic Local Alignment Search Too) [40] with the National Center for Biotechnology Information RefSeq (refseq_rna) as the reference database [41]. Alpha diversity was calculated using the inverse Simpson index at the OTU level [42].

#### Microbial Predictors

We analyzed obligate anaerobic bacteria by major anaerobic groups defined at various taxonomic levels for their importance in maintaining ‘healthy’ immunity [35, 43-46]: *Clostridia* (class), *Bacterioidetes* (phylum), *Negativicutes* (class) and *Fusobacteria* (genus). Percent anaerobes in a given stool sample was calculated by adding the percent 16S rRNA gene sequences of these obligate anaerobic bacteria.

### Monocyte subsets, analysis of blood samples, monocyte isolation and flow cytometry

A median of 7 blood samples were obtained from each patient during transplant days −10 to +30. 5-10 cc of blood in heparinized tubes were processed within four hours of collection. The first blood sample was collected within two days of hospital admission (prior to any perturbations to white blood cells from chemotherapy or radiation) and subsequent blood samples were collected at the start of white blood cell reconstitution through engraftment [47, 48].

Peripheral blood mononuclear cells were isolated by density gradient centrifugation (Histopaque 1119; Sigma). Single-cell suspensions were stained for surface antigens with fluorescently conjugated antibodies and samples were acquired with LSR II (Becton Dickinson). All flow cytometry data were analyzed using FlowJo software. For flow cytometry staining, the following antibodies were used: CD14-PE (clone M5E2; BD Biosciences), CD16-FITC (clone 3G8;BD Biosciences), CCR2-APC (clone K036C2; BioLegend), CD45-Alexa Fluor 700 (clone Hl30; BioLegend), CD11b (APC-Cy7; clone lCRF44; BioLegend), HLA-DR-PE-Texas Red (clone L243; BioLegend), CD86-PE-Cy7 (clone lT2.2; BioLegend), CD15-Pacific Blue (clone Hl98; eBioscience), CD20-PerCP-Cy5.5 (clone 2H7; BioLegend), CD3-PerCP-Cy5.5 (clone OKT3; BioLegend), CD19-PerCP-Cy5.5 (clone SJ25C1; BioLegend), CD56-PerCP-Cy5.5 (clone 5.1H11; BioLegend), and CD5-PerCP-Cy5.5 (clone UCHT2; BioLegend). Fluorescent minus one controls (FMO) were used to determine positive staining gate [49]. The monocyte gating strategy employed to define monocyte subsets (classical, intermediate, and non-classical) [10, 11, 50] is shown with CD14 on the x-axis and CD16 on the y-axis. A dump gate excluded B cells, T cells, and NK cells using CD19/CD20, CD3, and CD56, respectively **(Supplementary Figure 2)**.

### Analytical Approach

We analyzed absolute monocyte count recovery as a function of time at the start of immune reconstitution defined as: engraftment day minus 2 days (“reconstitution day”) until hospital discharge, using conditioning intensity and transplant type as stratification variables. Linear mixed models were fit using patient as a random effect (for intercept), absolute monocyte count as the dependent variable, and reconstitution day as a fixed effect. Fixed effect sizes and 95% confidence intervals are shown.

Descriptive statistics (obox plots) were used to visualize the proportions of classical (CD14^hi^CD16^neg^), intermediate (CD14^hi^CD16^Int^), and non-classical (CD14^Int^CD16^hi^) monocytes in the last blood sample collected from each patient, to assess whether the subpopulations of circulating monocytes reached their estimated targets: ∼85% classical monocytes; ∼10% non-classical monocytes; ∼5% intermediate monocytes. These thresholds were determined based on previous studies [10, 11, 51].

We calculated the Pearson correlation coefficient to measure the strength of the relationship between the proportion of anaerobic commensal gut microbes and microbial diversity to monocyte subset recovery. Next, we trained linear regression models while controlling for false discovery rates [52] to assess whether or not different clinical predictors correlated with monocyte recovery in the last blood sample tested (using the above mentioned ‘target’ values). The following clinical variables were used in the regression: 1. conditioning type (RIC versus MAC) 2. transplant type (T-cell depleted versus unmodified versus auto-HCT) 3. GCSF administration within 7 days of blood collection 4. A bloodstream infection (excluding positive blood cultures considered to be a “contaminant”) within 7 days of blood collection and 5. percent gram-negative bacteria (in the phyla, *Bacteroidetes* and *Proteobacteria*, major gram-negative taxa) in the stool sample collected within 3 days of the last blood sample collected. Statistical analyses were performed using R (v. 3.3.1).

## RESULTS

### Description of study population and biospecimens

Our cohort consisted of 18 patients who underwent auto- or allo-HCT at MSKCC between July 2015 to January 2016. Patients underwent different types of HCT for different hematologic malignancies. Four patients received an auto-HCT after myeloablative conditioning (MAC) with Carmustine (BCNU), etoposide, cytarabine, and melphalan (BEAM). Four patients received an allo-HCT after MAC conditioning consisted of the following regimens: total body irradiation (TBI), thiotepa, and cyclophosphamide or busulfan, melphalan, and fludarabine. Ten patients were recipients of an allo-HCT after reduced intensity conditioning (RIC) regimens. Clinical characteristics for each patient are shown in Table 1.

The duration of transplant hospitalization ranged from 20 to 38 days, during which antibiotics were given for both prophylactic and treatment purposes. Throughout this period, we sought to collect fecal samples on a daily basis. Out of the total 352 hospital days for these 18 patients, 318 stool samples were collected (90% of total hospital days). Of those samples, 236 (74%) yielded 16S amplicons that could be sequenced. Each blood sample was paired with a stool sample collected within 3 days. A median of 15 stool samples and 7 blood samples were collected per patient during their HCT hospitalization **(Table 1)**.

### Monocyte Recovery

Monocyte counts were considered during an analysis window from two days prior to the date of neutrophil engraftment (“reconstitution day”) until hospital discharge. Significant differences in reconstitution trajectories were found among groups stratified by conditioning regimens and transplant type. Patients receiving reduced intensity conditioning and unmodified transplants demonstrated the most variability (CI_95%_ = [0.196±0.062] and CI_95%_ = [0.194±0.060], respectively). All 18 patients reached a minimum normal absolute monocyte count value of 0.26 × 10^9^/L defined elsewhere, [53-55] and patients who received RIC and an unmodified transplant were more likely to overshoot the upper reference limit of 1.3 × 10^9^ (a value determined by the MSKCC laboratory) **(Figure 1A, Figure 1B)**.

**Figure 1.**
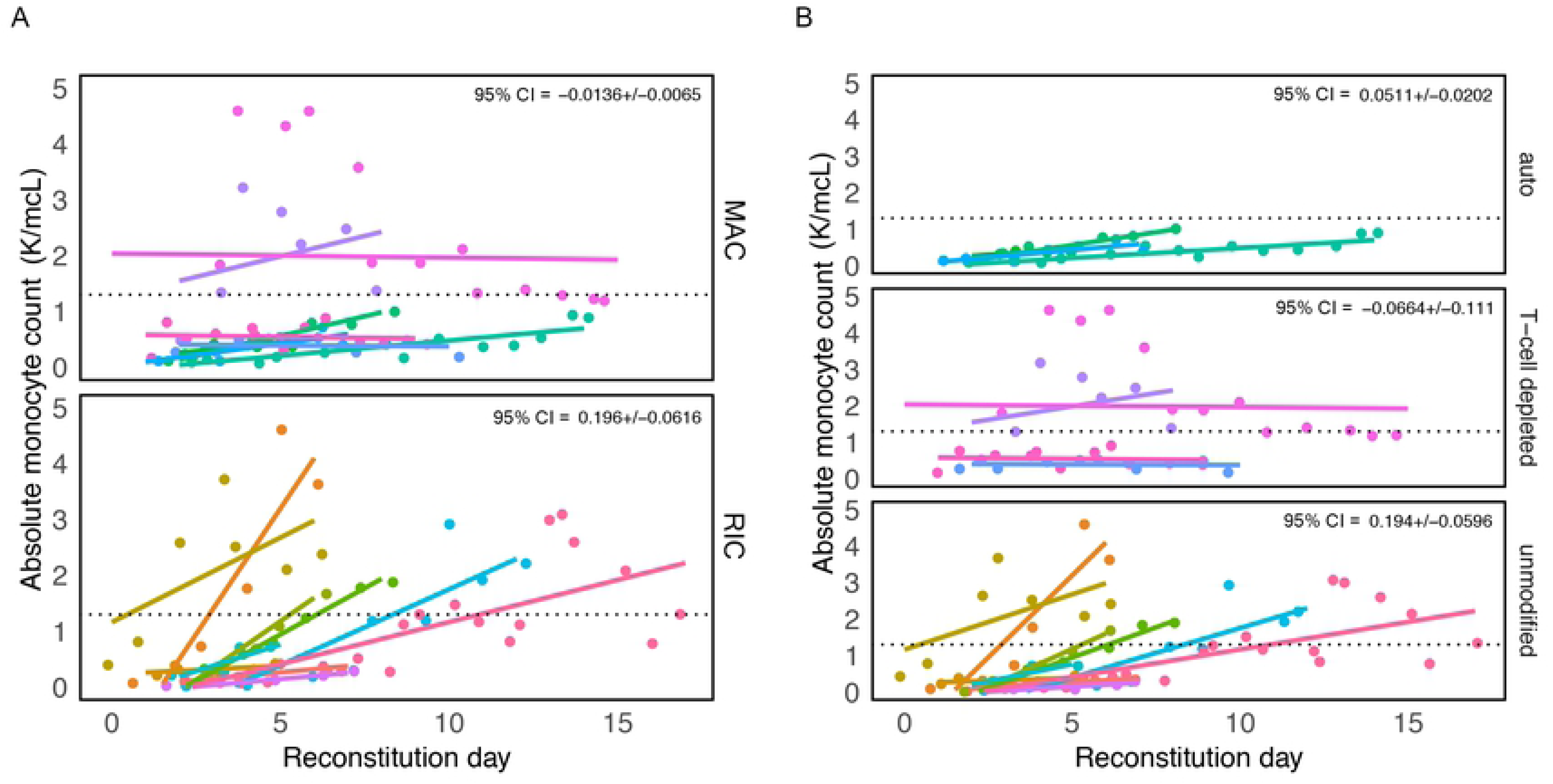
Recovery of absolute monocyte counts (AMC) is variable among patients receiving different conditioning regimens and transplant types. Graph showing absolute monocyte counts obtained for all patients when there was any sign of white blood cell recovery defined as “reconstitution day” (engraftment day – 2). Patient groups were divided by **(A)** conditioning, reduced intensity conditioning (RIC) and myeloablative conditioning (MAC) and **(B)** transplant type. Each color represents a different patient and a dashed line in each panel marks the upper limit of a “normal” AMC, defined by the MSKCC laboratory to be 1.3 K/mcL. In the upper righthand corner is the coefficient (i.e. slope/effect size) multiplied by 1.96 (+/- 95% confidence interval).

Three representative flow cytometry plots collected longitudinally from 3 patients who received different transplant types (Auto-HCT; Allo-HCT (RIC); Allo-HCT (MAC) show significant monocyte heterogeneity. That is, the commonly observed distribution of monocytes (‘banana shape’) within a patient [56, 57] changes over time **(Figure 2A)**. All other patients (n = 15) are shown in **Supplementary Figure 1.** The gating strategy used to identify the monocyte subsets of interest is shown in **Supplementary Figure 2**.

**Figure 2.**
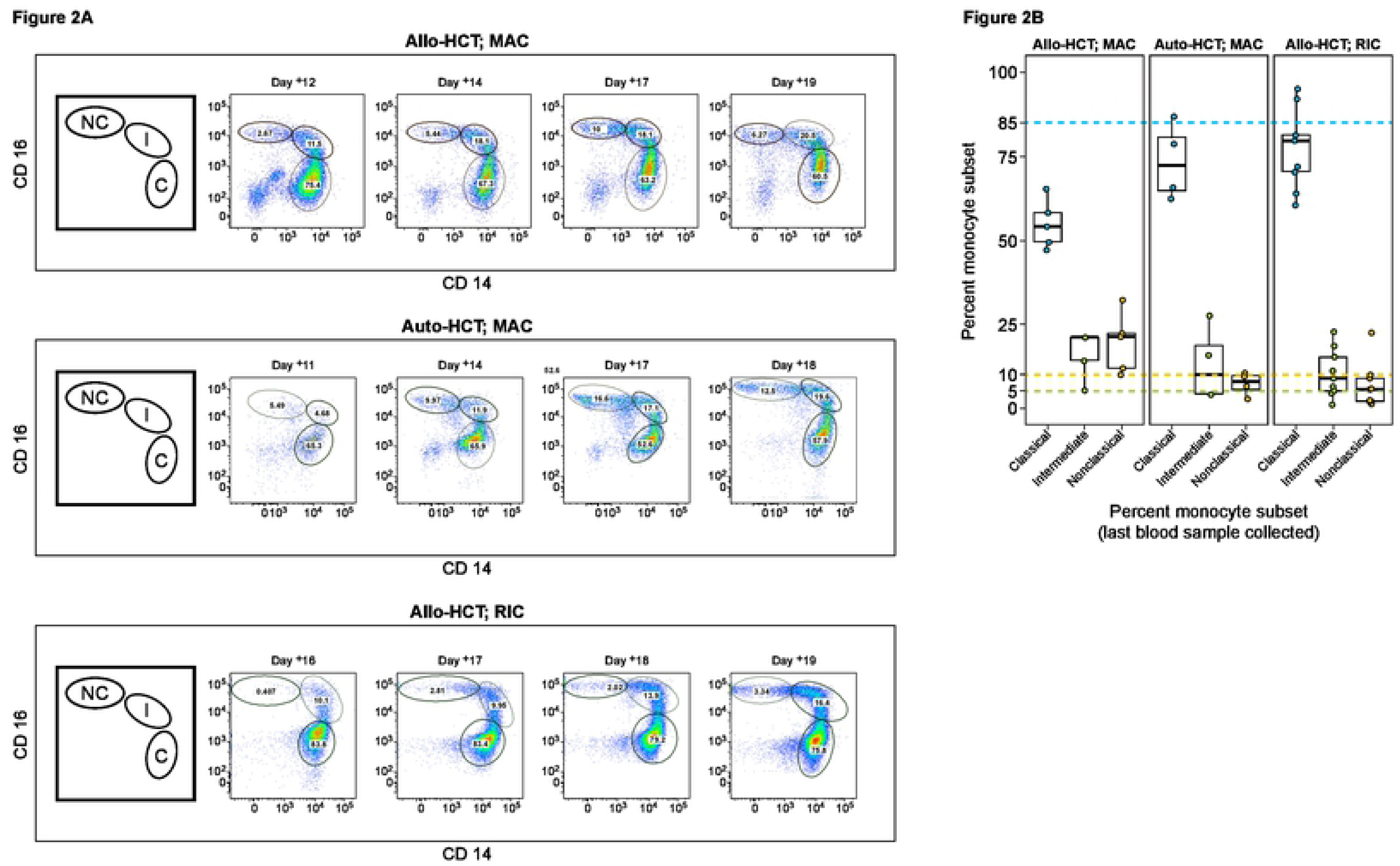
Variable recoveries of monocyte populations (classical, intermediate, and non-classical). **(A)** Time series for 3 representative patients using transplant day: Auto-HCT (top panel), Allo-HCT (reduced intensity conditioning (RIC; middle panel), and Allo-HCT myeloablative conditioning (MAC; bottom panel). The first figure (from left-to-right for each panel) is a cartoon illustration depicting each monocyte subtype, classical (C), intermediate (I), and non-classical (NC) (clockwise from left to right) that corresponds to successive multicolor flow cytometry plots that follow. The numbers in each gate are percentages of each monocyte population. The gating strategy is detailed in Supplementary Figure 1. **(B)** A boxplot showing the distribution of monocyte subsets determined from a patient’s last blood sample collected. Colored dashed lines indicate the upper limit of “normal” for the percent values for each monocyte subset: 85% classical monocytes (blue), 10% non-classical monocytes (yellow), 5% intermediate monocytes (green). Data points (dots) indicate the monocyte subset type using the same color scheme.

Some patterns in monocyte reconstitution were present; classical monocytes had an initial robust recovery consistent with previous findings [58], followed by restoration of intermediate and non-classical monocytes at later timepoints **(Figure 2A)**. This observation was especially true with respect to the recovery of non-classical monocytes. On average, patients who were recipients of either an auto-HCT or allo-HCT with RIC were more likely than allo-HCT patients with MAC to recover classical monocytes to target threshold (∼85%). The threshold of ∼5% intermediate monocytes was met in 72% (13/18) of patients, while those that received an allo-HCT with MAC conditioning were more likely to be discharged with levels of circulating non-classical monocytes that met target (∼10%) **(Figure 2B)**.

We next compared the frequency of circulating monocytes subsets with the composition of the intestinal microbiota. Diversity of intestinal microbial communities, as measured by inverse Simpson index, were not associated with monocyte recovery **(Figure 3A)**. Additionally, the proportion of obligate anaerobes associated with a healthy flora (*Negativacutes, Clostridia, Bacteroidiales*, and *Fusobacteria*) within the microbiota did not correlate with monocyte recovery **(Figure 3B)**.

**Figure 3:**
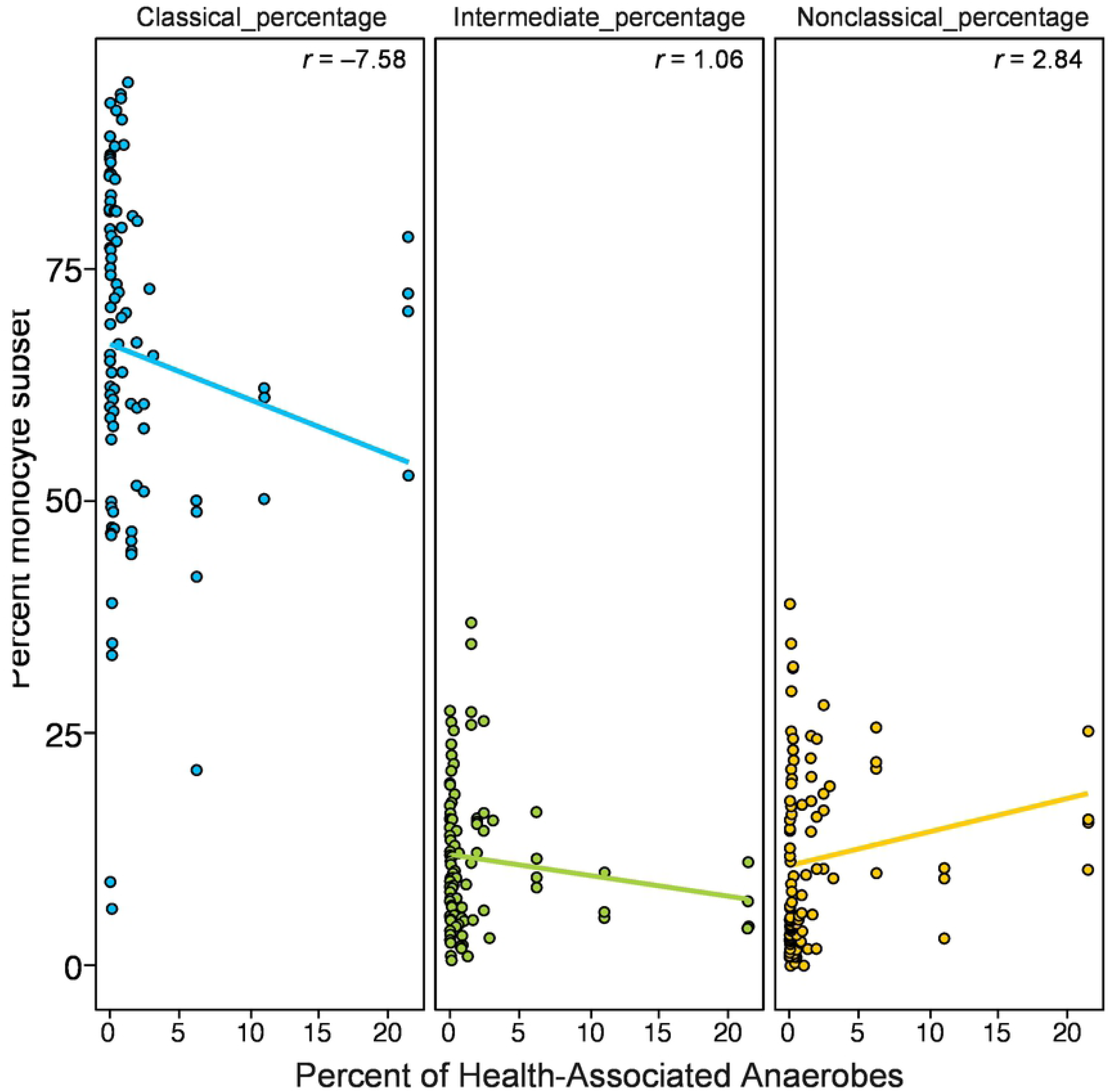
Monocyte subset frequency does not correlate with microbiota diversity or proportion of obligate anaerobes. **(A)** Relationship between the proportion of each monocyte subset and microbiota diversity measured by Inverse Simpson and **(B)** the proportion of obligate anaerobes (percent 16S rRNA gene sequences of *Negativacutes + Clostridia + Bacteroidiales +Fusobacteria*) in the stool microbiota for each matched stool and blood collection. Pearson correlation coefficient values are shown in the upper righthand corner of each panel.

While the proportion of monocyte subset recovery did not correlate with the microbiota, we next sought to investigate whether immune activation status of monocytes subsets was altered by the microbiota composition. Monocyte immune activation phenotype as measured by the expression of cell surface markers (CD86, HLA-DR, and CCR-2 **(Supplementary Table 1)**). was assessed in each three subsets of monocytes. No correlation was observed between expression of these markers and microbiota diversity **(Supplementary Figure 3)** or percent of obligate anaerobic bacteria in the microbiota **(Supplementary Figure 4)**.

We used a simple linear regression model to test whether different compositional characteristics of the microbiota and different clinical variables are related to monocyte recovery. In a linear regression model, microbiota diversity and the proportion of gram-negative Proteobacteria? in the gut were not observed to be associated with monocyte recovery of each subset. Exposure to GCSF within 7 days of blood sample collection/processing was associated with successful reconstitution of classical and intermediate monocytes (p = 0.037 and p =0.002, respectively). Conversely, a T-cell depleted transplant was negatively associated with these monocyte subsets (p = 0.041 and p = 0.008, respectively) **Table 2**. Non-classical monocytes were not associated with any of the clinical predictors we defined.

## DISCUSSION

HCT involves the administration of intense chemotherapy with or without radiation and antibiotic regimens, resulting in large shifts in leukocyte and microbiota compartments, often leading to complete compositional changes from one day to the next. The myeloablation associated with HCT provides a unique platform for exploring the nascent, re-development of immune reconstitution. Prior studies have supported the symbiotic relationship between the gut microbiota and the systemic immune system [59, 60], but most prior work has focused on the role the microbiota has on shaping different populations of T cells [30-32]; little is known about the impact intestinal commensalism has on monocyte maturation. We followed a systematic strategy to identify three subsets of monocytes (classical, intermediate, and non-classical monocytes) [10] during immune reconstitution post-HCT to assess whether constituents of the microbiota affect monocyte recovery.

This pilot study involved high-frequency collection of stool and blood samples from 18 patients to assess whether a link between the microbiota and monocyte recovery could be found. We found no correlation between the microbiota composition and differences in monocyte recovery when analyzing a higher-diversity microbiota or a microbiota composed primarily of either commensal anaerobes or Gram-negative organisms. We also found no association between microbiota composition and the expression of co-stimulatory markers, including CD86, HLA-DR and the chemokine receptor, CCR2.

Analysis of factors associated with immune reconstitution revealed that any link between the microbiota and immune reconstitution is weak relative to other clinical variables. As expected, GCSF exposure correlated positively with classical and intermediate monocyte reconstitution, while T cell depletion by means of CD34 positive selection was negatively associated with classical and intermediate monocyte subset reconstitution. We also found significant variability in the reconstitution of absolute monocyte counts and monocyte subsets across patients. The biological significance that the varying rate of monocyte recovery has on patient outcomes needs to be further explored.

Discerning the effects of many patient variables in the setting of HCT, including medications delivered and clinical complications that ensue (bacteremia, fever, GVHD, etc.) make forming a direct relationship between the microbiota and monocytes in HCT patients challenging [61]. Medications given to HCT patients that have immunomodulating properties (i.e., antibiotics and steroids), could serve as confounders in our analysis. For example, 16/18 patients received ciprofloxacin and 7/18 patients received steroids. Ciprofloxacin can dampen the effects of LPS [62, 63], a potent stimulator of monocyte mobilization out of the bone marrow [30, 64], while steroids can reduce blood levels of monocytes [30, 65]. Further, testing for intestinal LPS permeability in human serum or blood is technically challenging [61].

While we were not able to discern an association between microbiota composition and monocyte recovery in this heterogeneous pilot cohort, this study demonstrates the feasibility of high-resolution sampling of blood and stool samples and opens the door to future studies where sampling many more patients with daily frequency may help distinguish which microbial and clinical factors drive HCT patient outcomes.

## Figure Legends

**Supplementary Figure 1: Circulating monocyte subsets in transplant patients following immune reconstitution.** Time series of blood samples collected for 15 representative patients with transplant day used to show when samples were collected: Auto-HCT patients (top panel), Allo-HCT patients (RIC) (middle panel), and Allo-HCT patients (MAC) (bottom panel). A cartoon illustration in the upper righthand corner of each panel is again shown depicting the relative location of each monocyte subtype, classical (C), intermediate (I), and non-classical (NC) (clockwise from left to right).

**Supplementary Figure 2: Monocyte gating strategy. (A)** Cells are first visualized on Live/Dead vs. FSC and a gate is drawn around the live cells. **(B)** A gate is drawn around lymphocytes and monocytes (SSC vs. FSC). **(C & D)** Doublets are then discriminated in two steps (FSC-A VS. FSC-W and SSC-A and SSC-W). **(E)** CD45+ (hematopoietic cells), lineage negative (Non-T, Non-B, Non-NK) cells were gated for. **(F)** Gating to further discriminate monocytes from granulocytes (CD15+, CD16+). **(G)** (CD14+CD16+) monocytes subsets (classical, intermediate, non-classical) are shown (plot f). Numbers are percentages of each population within each gate.

**Supplementary Figure 3: Monocyte expression of immune activation markers does not correlate with microbiota diversity.** Relationship between compositional diversity of the gut microbiota, measured by Inverse Simpson and the proportion of surface marker expression of each monocyte subsets (classical, intermediate, and nonclassical): CCR2 (top panel), CD86 (middle panel) and HLA-DR (bottom panel). Pearson correlation coefficient values are shown in the upper righthand corner of each panel.

**Supplementary Figure 4: Monocyte expression of immune activation markers does not correlate with proportion of obligate anaerobes in the microbiota.**

Relationship between the proportion of commensal anaerobes that make up the gut microbiota and the level of surface marker expression of each monocyte subsets (classical, intermediate, and nonclassical): CCR2 (top panel), CD86 (middle panel) and HLA-DR (bottom panel). Pearson correlation coefficient values are shown in the upper righthand corner of each panel.

**Table 1: Clinical characteristics of all 18 HCT patients including the number of blood samples and stools samples collected from each patient.** Abbreviations: Pt, patient; Auto, autologous; Allo, allogeneic; RIC, reduced intensity conditioning; MAC, myeloablative conditioning; DLBCL, diffuse large B cell lymphoma; MDS, myelodysplastic syndrome; CLL, chronic lymphocytic leukemia; ALL, acute lymphoblastic leukemia; AML, acute myeloid leukemia; cy, cyclophosphamide.

**Table 2: Parameters assessed for association with monocyte subset recovery** Abbreviations: Auto, autologous; Allo, allogeneic; RIC, reduced intensity conditioning; MAC, myeloablative conditioning; GCSF, granulocyte colony-stimulating factor.

**Supplemental Table 1:**

Fluorochromes used to label antibodies used and their function

## Footnotes

### Meeting(s) where the information has previously been presented

The information contained in this paper has not been presented at previous meetings

